# Assessing mechanical agency during apical apoptotic cell extrusion

**DOI:** 10.1101/2023.10.26.564227

**Authors:** Sommer Anjum, Llaran Turner, Youmna Atieh, George T. Eisenhoffer, Lance Davidson

**Author notes:** authors for correspondence: Lance Davidson,; George Eisenhoffer. lead contacts: Lance Davidson,; George Eisenhoffer.

## Abstract

Epithelial tissues maintain homeostasis through the continual addition and removal of cells. Homeostasis is necessary for epithelia to maintain barrier function and prevent the accumulation of defective cells. Unfit, excess, and dying cells can be removed from epithelia by the process of extrusion. Controlled cell death and extrusion in the epithelium of the larval zebrafish tail fin coincides with oscillation of cell area, both in the extruding cell and its neighbors. Both cell-autonomous and non-autonomous factors have been proposed to contribute to extrusion but have been challenging to test by experimental approaches. Here we develop a dynamic cell-based biophysical model that recapitulates the process of oscillatory cell extrusion to test and compare the relative contributions of these factors. Our model incorporates the mechanical properties of individual epithelial cells in a two-dimensional simulation as repelling active particles. The area of cells destined to extrude oscillates with varying durations or amplitudes, decreasing their mechanical contribution to the epithelium and surrendering their space to surrounding cells. Quantitative variations in cell shape and size during extrusion are visualized by a hybrid weighted Voronoi tessellation technique that renders individual cell mechanical properties directly into an epithelial sheet. To explore the role of autonomous and non-autonomous mechanics, we vary the biophysical properties and behaviors of extruding cells and neighbors such as the period and amplitude of repulsive forces, cell density, and tissue viscosity. Our data suggest that cell autonomous processes are major contributors to the dynamics of extrusion, with the mechanical microenvironment providing a less pronounced contribution. Our computational model based on *in vivo* data serves as a tool to provide insights into the cellular dynamics and localized changes in mechanics that promote elimination of unwanted cells from epithelia during homeostatic tissue maintenance.

## INTRODUCTION

Cell extrusion is critical for the maintenance of epithelial tissues (Rosenblatt et al., 2001), as dysregulation of cell extrusion can disrupt epithelial barrier function and drive formation of adenocarcinomas (Eisenhoffer and Rosenblatt, 2013). Key modes of homeostatic cell extrusion include apoptotic cell extrusion, extrusion of transformed cells, and extrusion of live cells after overcrowding (Eisenhoffer et al., 2012; Katoh and Fujita, 2012; Villars and Levayer, 2022). An inducible *in vivo* experimental system for apoptotic cell extrusion was recently developed in epithelium of the larval zebrafish tail fin (Atieh et al., 2021b). This epidermis is best classified as a squamous-type epithelium. Transgenic fish express the genetically encoded enzyme nitroreductase (NTR) in a subset of cells of the larval epidermis and addition of the prodrug metronidazole (MTZ) creates a cytotoxic byproduct that promotes cells to be extruded from the epidermis. We collected high resolution time-lapse sequences of extruding epidermal cells and quantified geometric parameters of both extruding cells and their neighbors; before, during, and after extrusion. Cells in NTR/MTZ-activated epithelia undergo a series of actomyosin-driven pulsatile contractions that result in oscillations of cell area that can be characterized by amplitude and duration. Area oscillations in extruding cells are additionally regulated by sphingosine-1-phosphate (S1P) and activation of a caspase cascade. Similar regulation of apoptotic apical extrusion involving actomyosin contractility by S1P and RhoA have been observed in cultured epithelial cell lines (Duszyc et al., 2021; Villars and Levayer, 2022)

Oscillations in the shape of apoptotic extruding cells raise several questions concerning the mechanical agency of extruding cells and the dependence of extrusion on the microenvironment. What is the role of the local mechanical environment in promoting extrusion? Epithelial cell behaviors are commonly regulated and constrained by their neighbors. Are the pulsatile contractions coordinated between cells? Epithelial cells in many embryos and larvae exhibit area oscillations, but it is unknown if these oscillations are adaptive. Does a cell facilitate its own exit or is it pushed? Extrusion involves dynamic changes in cell adhesion and contractility in both the extruding cell and its neighbors, but we know little about the relative mechanical contributions of these cell biological changes. To address these questions, we divide our interest in the biophysics of extrusion into two categories, first, considering processes operating in the extruding cell (e.g. autonomous mechanical processes), and second, processes modulating extrusion that operate within cells adjacent to the extruding cell (non-autonomous mechanical processes). Akin to intrinsic properties, autonomous mechanics refers to the biophysical properties of the extruding cell that influence the kinematics of extrusion, including steady state mechanical properties as well as the strength, rate, and periodicity of cyclic contractility. Conversely non-autonomous processes refer to properties of the extruding cell’s microenvironment such as the neighboring cells’ mechanical properties and cyclic contractility, and tissue confinement. It is currently not possible to precisely manipulate biophysical properties such as amplitude, frequency, or packing density of live extruding cells, or their microenvironment. To address these outstanding questions, we sought to build a computational model where the roles of cell-autonomous and non-cell-autonomous mechanics in driving apical extrusion could be tested and compared quantitatively.

Epithelial homeostasis and morphogenesis is commonly simulated within one of four computational frameworks (Davidson et al., 2010; Fletcher et al., 2017). The most simple physical analog of extrusion is the so-called T2 transition within soap film arrays (e.g. wet foams), where cell loss occurs through a 3-neighbor cell array (Weaire and Hutzler, 2001). Most epithelial tissue models assume a linearly elastic tissue that is largely homogeneous and stable. Vertex models are the most common biophysical models of epithelia (Staple et al., 2010). In vertex models, the dynamics of apical epithelial vertex movements minimize area and perimeter deviations from preferred values amidst changing force balances between the medioapical cell cortex and apical cell-cell junctions (Farhadifar et al., 2007; Fletcher et al., 2014; Hashimoto et al., 2015; Honda et al., 2008; Rauzi et al., 2008; Spahn and Reuter, 2013; Weliky et al., 1991; Weliky and Oster, 1990). A vertex model of delamination from an overcrowded tissue (Marinari et al., 2012a) incorporated triangular cells with small areas that can be removed. However, the dynamics of extrusion in this work are not explicitly defined and the removal is not encoded in the system. Another vertex model instead interrogated model parameters needed to form a rosette characteristic of the extrusion process through significant increases of extruding cell contractility and local adhesions (Kuipers et al., 2014). A Cellular Potts Model also facilitates simulation of epithelial cell dynamics, where packets of cytoplasm are distributed among cells in an epithelium to minimize an energy functional representing apical and junctional elastic energy, as well as relevant phenomenological processes (Belmonte et al., 2016; Graner and Glazier, 1992; Hirashima et al., 2017; Wolff et al., 2019). Finite element models can represent epithelial tissue-scale mechanics but can also include substructures such as protrusions and cell junctions, and superstructures such as multicellular contractile belts and swelling extracellular matrix (Brodland et al., 2009; Chen and Brodland, 2000; Davidson et al., 1999; Ramasubramanian and Taber, 2008). Agent-based, or active particle models represent epithelial cells, or their centers, as particles that can be endowed with biophysical properties and rules governing their interactions (Dalle Nogare and Chitnis, 2020). Cell junctions composing an epithelial sheet in an active particle model are represented by tiling or tessellating a set of cell centers (Bi et al., 2015; Bi et al., 2016). Of these four types of models, active particle models can be developed with minimal sets of parameters and implemented with a limited set of assumptions on cell adhesions and intracellular remodeling.(Manning and Collins, 2015). Each of these simulation frameworks can represent discrete cells and enable direct quantitative comparison with experiments. As of yet, no computational models have sought to represent oscillatory extrusion and the contribution of the local mechanical microenvironment.

To explore the biophysical principles regulating cells extrusion in the larval zebrafish, we developed an active particle model that represents the dynamic and stochastic mechanics of the extruding cell and its surroundings. Such a model needed to account for several experimental observations including a sudden drop in area upon the onset of extrusion, and rachet-like oscillatory decreases in area in the extruding cell with a range of amplitudes and periods. While active particle models have been useful in the analysis of epithelial morphogenesis and solid-to fluid-like transitions that allow jammed tissues to flow, these models have not previously been used to investigate extrusion. Initially, we sought to represent apical cell junctions by tiling cell centers via Voronoi tessellation. However, we found such tessellation schemes could not represent mechanically heterogenous cell populations, but only the geometry of cell positions (Aurenhammer and Edelsbrunner, 1984; Du et al., 1999; Sánchez-Gutiérrez et al., 2016). To properly represent both the mechanical and geometrical heterogeneity of cells in a 2D sheet, we developed a hybrid tessellation that integrated mechanical interactions within the centroidal Voronoi Tessellation framework (hybrid CVT). This hybrid tessellation framework allowed extruding cells to be represented with small areas and mechanically driven oscillations. We simulate mechanical interactions through repulsive cell potentials and incorporate these potentials to map cell-cell boundaries of extruding cells and their neighbors. In addition to providing control over cell-to-cell biophysical properties, the model allows us to test the effect of extrusion via tissue-wide properties such as viscosity and cell packing. By testing each of these features, we have been able to quantify the role of extruding cell autonomy and demonstrate that nonautonomous factors may play a smaller role than earlier suggested.

## RESULTS

### Topology of the zebrafish larval epidermis

To better understand variation of the size and shape of cells within the zebrafish larval epidermis, we first determined the polygon distribution of cells within the epithelium under homeostatic conditions and captured how these parameters could fluctuate from larvae to larvae. Confocal images of the tail fin epithelium of transgenic larvae were collected 4 days post-fertilization (dpf) were collected using fluorescent proteins that localize to epithelial cell-cell junctions (cldnB::lyn-GFP) and nitroreductase as part of the inducible apoptotic system(NTR-mCherry) (Figure 1a). We confirmed the expression of with a closer look at the larval zebrafish tail fin (Figure 1b). Morphometric analysis showed over 55.7% of cells in the epithelium have an area of 800-1000µm^2^ in a skewed-right distribution (Figure 1c), and are comprised of 29.9% pentagons, 34.2% hexagons, 13.8% heptagons, 1.9% octagons and 0.2% nonagons (Figure 1d). These data are consistent with the observed polygon topology of the epidermis in other eukaryotes, including the epidermis of the *Xenopus* tadpole tail, the outer epidermis of the freshwater cnidarian *Hydra,* the *Drosophila* wing disc epithelia, and the epidermis of plants (Gibson et al., 2006). In all of these cases, a similar non-gaussian distribution of epithelial polygons is observed with less than 50% hexagonal cells and high percentages of pentagonal and heptagonal cells. These data support the idea that similar topological distributions are present in other multicellular eukaryotes and provide the basis for a ground state for a computational model of epithelial dynamics.

**Fig 1.**
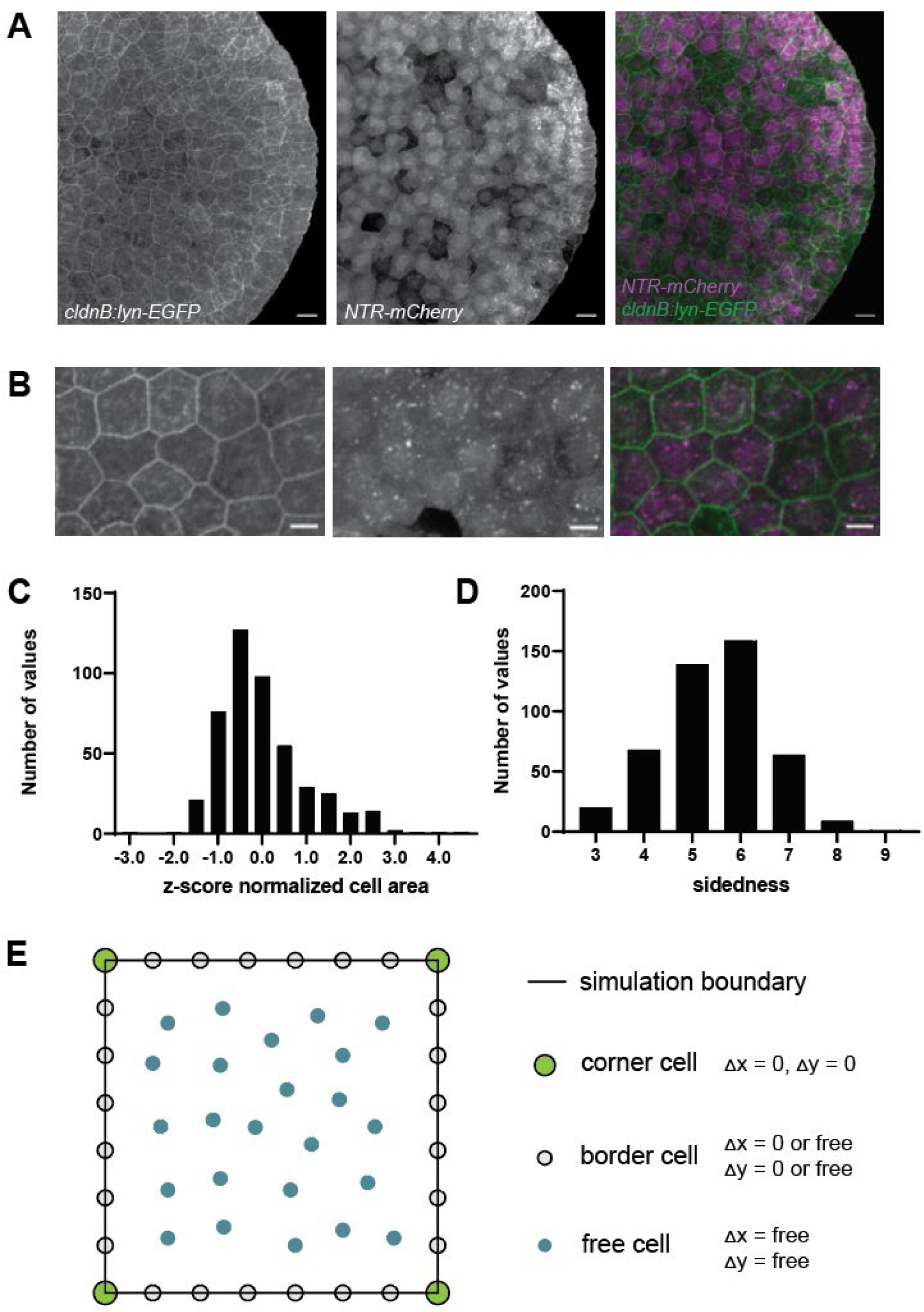
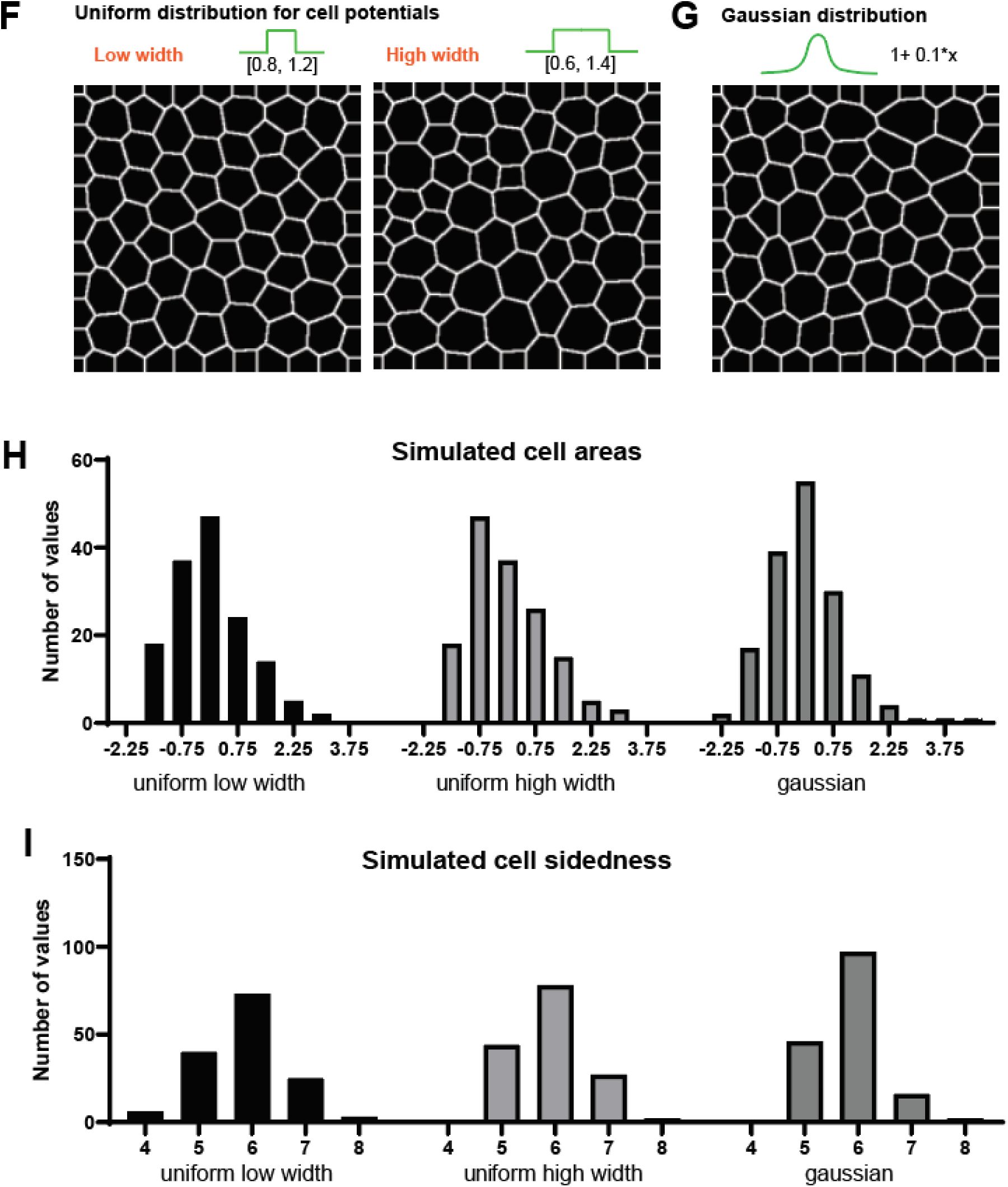
In vivo and in-silico epithelium at equilibrium. (a) The larval zebrafish tail fin epithelium with cell-cell junctions fluorescently labeled. (Scale bar = 20μm) (b) Close up of larval zebrafish epithelium with the expression of NTR-mCherry (Scale bar = 10μm) (c) Z-score normalized distribution of areas of tailbud epithelium cells (d) Sidedness distribution of tailbud epithelium cells (n= 10 larvae, 464 total cells) (e) Schematic of types of cells in simulation and their movement restrictions (f) Cell fields obtained from using low and high width uniform distributions for sampling cell size (g) Cell fields obtained from using a Gaussian distribution for sampling cell size (h) Z-score normalized cell area distributions for the three distributions over 10 simulations each (n=141,151,161) (i) Cell sidedness distributions for the three distributions over 10 simulations each (n=141,151,161)

### Active particle representation of a mechanically and geometrically heterogeneous epithelium

To simulate an epithelial sheet of cells, we developed a lattice-free model where cells are represented by interacting particles, or nodes, that are positioned within a two-dimensional rectangular domain. Cells in this rectangular patch are represented by three types of nodes: 1) corner nodes represent quarter-cells that connect the four corners of tissue to the domain boundary, 2) border nodes represent half-cells that connect the four tissue edges to the domain boundary, 3) interior free nodes represent full sized cells in the epithelium within the domain. Corner nodes do not move, whereas border nodes are free to move along the axes that demarcate the domain boundary. Interior nodes may move freely within the 2D domain. Border and internal nodes move as they interact with each other and move towards mechanical equilibrium via repulsion forces operating between all nodes (Figure 1e). Node movements are governed by a force balance enforcing Langevin dynamics, incorporating stochasticity and drag forces from viscous media. We represent the mechanical elasticity of each cell with a non-linear repulsive spring. Each cell is a node with its own unique mechanical properties. We represent the mechanical heterogeneity of the epithelium by assigning each node a repulsive potential that is randomly selected from a uniformly distributed set of potentials/rest lengths (rest_i,j,k…_). Forces act within an interaction distance, *rest*, of cell *i* (Equation 1),

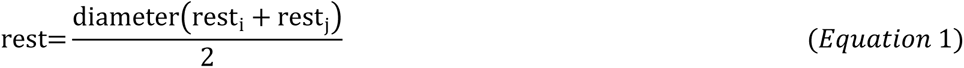

where *diameter* is an average cell diameter and rest_i_ and rest_j_ are rest lengths of cell *i* and other cells *j*. These rest lengths determine repulsion forces between two cells as a function of diameter and the distance between them (Equation 2),

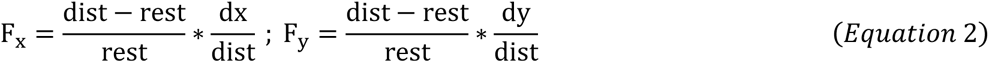

where the distance, *dist*, is calculated between adjacent nodes and resolved into its components *dx* and *dy* and used to calculate forces in the x and y directions, *F_x_* and *F_y_,* respectively. Force is calculated in the x and y directions between cell i and cells j at a distance less than *rest* away. The x and y forces are summed for all cells within the interaction distance (Equation 3),

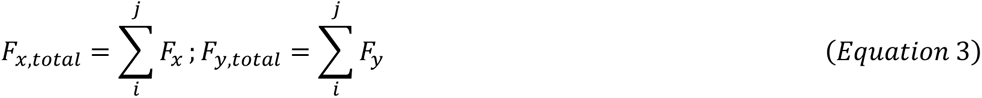

Cell motion occurs as force is scaled by viscosity and the simulation is advanced in time using forward Euler methods (Equation 4),

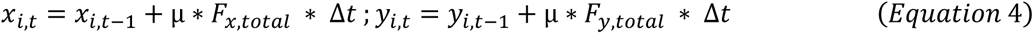

where each x and y position at a given time *t*, *x_i,t_* and *y_i,t_*, are calculated using the position in the previous time *t-1* and the calculated total force. Viscosity is represented by *μ* and change in time between *t* and *t-1* is *Δt*.

To visualize changes in epithelial cell morphology and network topology, we convert our array of nodes into a 2D graph of topologically connected cells with defined shapes and areas. One option for conversion is a Voronoi tessellation of the centroids. Voronoi tessellations are commonly used to mimic epithelial shape distributions in active particle models (Bi et al., 2016; Lin et al., 2014; Paszek and Weaver, 2004; van Drongelen et al., 2018). These tessellations have known limitations, particularly in representing cell shape asymmetries and heterogeneity present in tissues responding to anisotropic forces (Kaliman et al., 2016). A shortcoming of using the Voronoi tessellation with our study is that it cannot represent sufficiently small cells, even if cell centroids become close to each other (Figure S1a-d). One alternative is to use the centroidal Voronoi tessellation (CVT; see Supplement) to form the 2D graph; CVT determines the centroid of each triangle in a Delaunay triangulation and connects them with straight lines. However, this method is also purely geometric, like the Voronoi tessellation, and is equally incapable of representing the cell size heterogeneity present in our model system. In order for the geometric organization and arrangement of simulated cells to reflect the heterogeneity of cell sizes observed *in vivo*, we developed a new method. Like the CVT method, we first calculate the circumcenters of the triangle but then shift their positions by the weighted centroid of the potentials of the nearby cells, leveraging the physics of the system. We average the weighted centers with the original Voronoi circumcenters and connect them, creating our own hybrid CVT (Figure S1e). Our model, tessellation scheme, and analysis pipeline (supplement, Figure S2) enable us to recapitulate extrusion events in the zebrafish tail epidermis.

### Cell heterogeneity as captured by simulation

As there is variation in cell morphology from embryo to embryo, as well as across the same tissue, we developed our model to represent this variability. To generate diverse cell areas and sidedness in the epithelium at rest, we sample individual cell rest lengths from three distributions of rest lengths including two uniform distributions (Figure 1f) and an adjusted gaussian distribution (Figure 1g) centered at 1.0. Preliminary analysis indicated that tissue packing also affects area and sidedness, with lower packing yielding more varied area and sidedness, so we chose distributions that result in lower levels of packing (Figure S3). The area and sidedness across the three cases are not significantly different from each other (p>0.05; two-sample Kolmogorov-Smirnov test). Furthermore, area distributions are not significantly different from the area measurements from cell in living epithelia (Figure 1c,d), however, sidedness in the simulations differs from these *in vivo* data (Figure 1 h,i). To offset this difference, we chose to sample cell potentials from the uniform high width distribution as it yielded a lower sum of test statistics compared to the live cell area and sidedness (Table 1).

**Table 1.** Area and sidedness comparisons.

| Case | Area test statistic | Area p-value | Sidedness test statistic | Sidedness p-value | Total test statistic |
| --- | --- | --- | --- | --- | --- |
| Uniform low width | 0.10 | 0.18 | 0.18 | 1.20e-03 | 0.28 |
| Uniform high width | 0.08 | 0.51 | 0.20 | 1.48e-04 | 0.28 |
| Gaussian | 0.12 | 0.05 | 0.21 | 5.28e-05 | 0.33 |

### Cell potential oscillations generate kinematic changes in apical surface area and can be used to drive extrusion

In the larval zebrafish epithelium, cell areas vary in two distinct modes, decreasingly sinusoidal in extruding cells, and sinusoidally in neighboring cells (Figure 2a,c). We use area strain to compare changes in area over time (area_final_, area_initial_) to account for cell area variation:

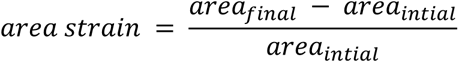

**Fig 2.**
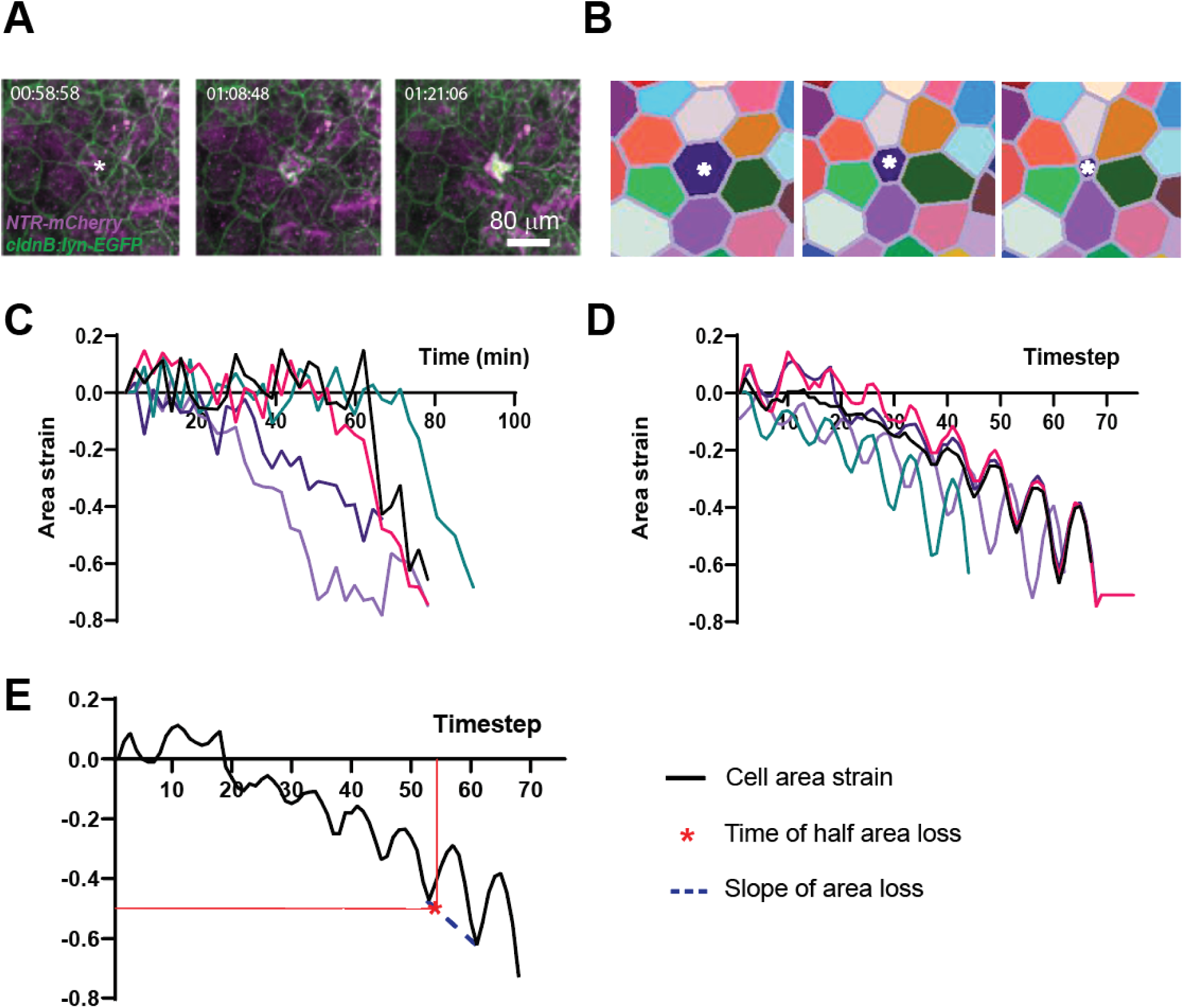
Quantifying and modeling epithelial cell extrusion. (a) Time-lapse imaging of cells extruding from the larval zebrafish epidermis (b) A simulated extruding cell over time (c) Five representative area strain trajectories of extruding cells from time-lapse imaging data (d) Five simulated cell area trajectories matching cell trajectories (e) Quantification of time to half area loss and slope of area loss

In order to drive cell area oscillation as seen *in vivo*, we apply a time-varying sinusoidal potential, rest_i_, to a cell (Equation 6),

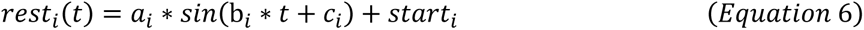

where ***a*** represents the oscillation amplitude, ***b*** represents the frequency, and ***c*** represents the phase of the oscillation.

The observed variations in cell areas within the larval zebrafish epidermis (Figure 1c, Figure 2c,e) may reflect cell-intrinsic mechanical differences. To represent this variation we assign different rest potentials to individual cells (start_i_, start_j_, start_k_). However, cell area fluctuation, or oscillation, is a common feature of animal epithelia, and *in vivo* observations of MTZ-treated tail epidermis reveal apical area oscillations during the process of cell elimination by extrusion. Therefore, oscillation through the sinusoidal term was added to the start potential (Equation 6). Two cells can oscillate in-phase, where high levels of potential are synchronized (c_i_ = c_j_ = 0 for both cells) or can oscillate out of phase (c_i_ - c_j_ ≠ 0°). In our model system, cell areas oscillate out of phase (Figure 2c,e). To make the oscillation cause cell area to decrease over time, we add a linear term with slope d_i_ (Equation 7),

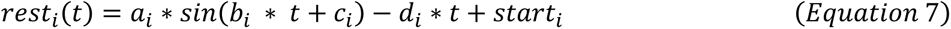

By implementing these equations, we can represent cell area changes seen *in vivo* during extrusion (Figure 2d). Thus, variations in cell area, and the dynamics of apical areas during extrusion can be regulated by a time-varying rest potential.

To quantify how changing the model parameters affects area strain in simulated extruding cells, we calculate the time to half area loss (t_half_) and the slope when the time to half area loss is reached (slope_half_; Figure 2e). Other endpoints such as local strain and strain along the x and y directions were considered but were not as varied across simulations (Figure S4).

### Base Extrusion Model: a minimal model of extrusion of an oscillating cell

Cell shape oscillation can be driven by varying the cell’s potential function over time. As individual cells are extruded apically after MTZ treatment, they exhibit a distinctive form of oscillation. The cell starts with an initial area that oscillates, and then the cell loses a large portion of its area (Figure 2c; Eqn. 7). Additionally, epidermal cells immediately adjacent to extruding cells also exhibit distinctive, high amplitude oscillations, also driven by a time-varying potential (Eqn. 6). While multiple cells can be made to extrude, we chose to have only one cell leave at a time to avoid effects of distant oscillations (Figure S5).

We sought to simulate key observations of extruding cells in living epithelia including: i) ratchet-like decreases in area followed by a sudden drop, ii) cells do not oscillate to areas greater than their initial area, and iii) amplitude and frequency of area change seen in cells via time-lapse imaging, and iv) area loss trend as seen *in vivo*. To accommodate this range of features, we varied parameters of the driving potential to mimic changes of experimentally measured cell areas. We ran simulations with different parameters and compared the extruding cell’s area trajectory to cell area trajectories observed in living epithelia. We compared the amplitudes of the extruding cells’ area trajectories between the *in vivo* and *in silico* cases, showing that they are significantly similar (Mann Whitney U, p>0.05, n=14 live cells, n=15 simulated cells). We selected a set of parameters that generated extrusion over several simulations matching cell data observed *in vivo* (Table 2). This parameter set represents a base case for the perturbations to follow.

**Table 2.**
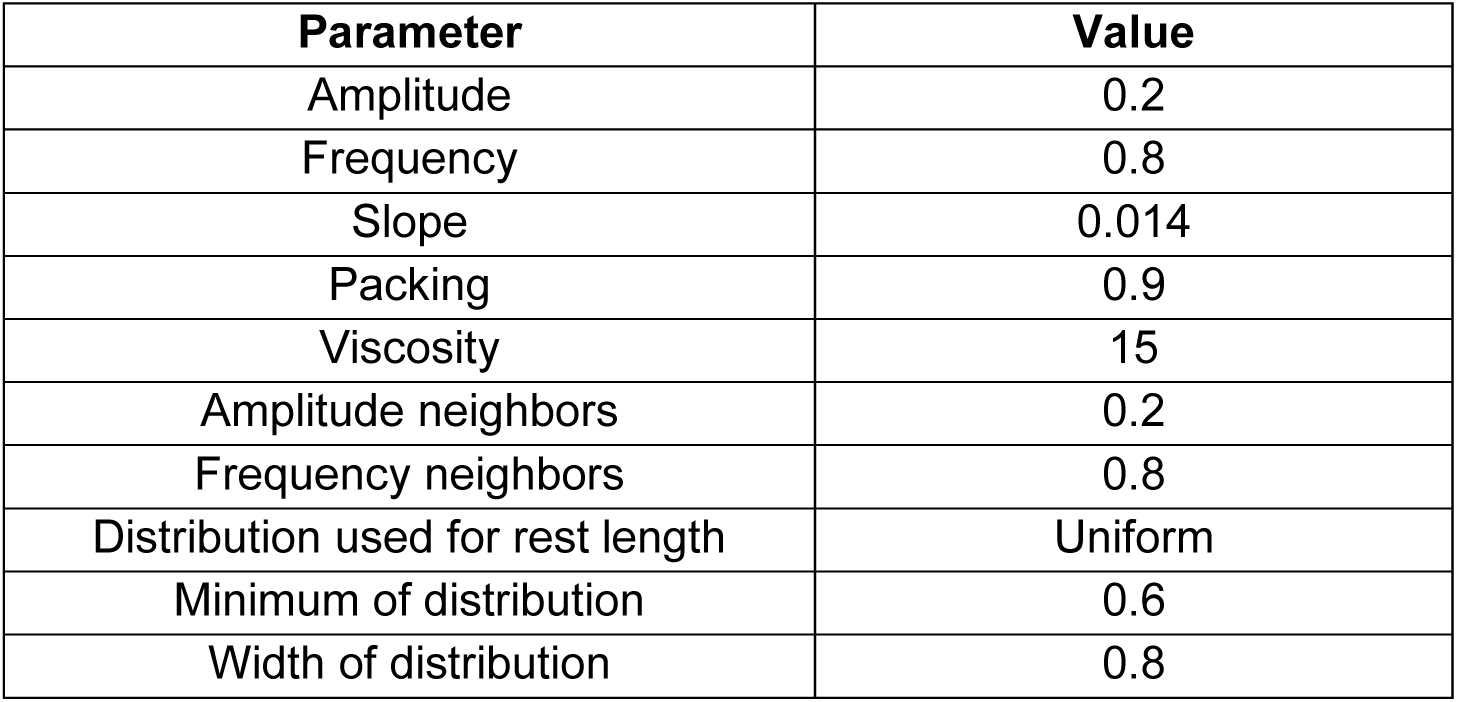
Parameter values for base case.

| Parameter | Value |
| --- | --- |
| Amplitude | 0.2 |
| Frequency | 0.8 |
| Slope | 0.014 |
| Packing | 0.9 |
| Viscosity | 15 |
| Amplitude neighbors | 0.2 |
| Frequency neighbors | 0.8 |
| Distribution used for rest length | Uniform |
| Minimum of distribution | 0.6 |
| Width of distribution | 0.8 |

Model parameters can be varied to assess their effect on t_half_ and slope_half_ (Figure 3). Within the cell autonomous regime, we can vary properties of the extruding cell potential, such as amplitude (*a*), frequency (*b*), and initial rest length (*start*). Beyond the extruding cell, we consider two nonautonomous contributors: the microenvironment and the macroenvironment. We can assess the effect of the microenvironment on extrusion by changing the neighbor cell potential function’s amplitude *(a*), frequency (*b*), and initial rest length (*start*). We can assess the role of the macroenvironment on extrusion by changing viscosity (*μ*) and packing (*p*).

**Fig 3.**
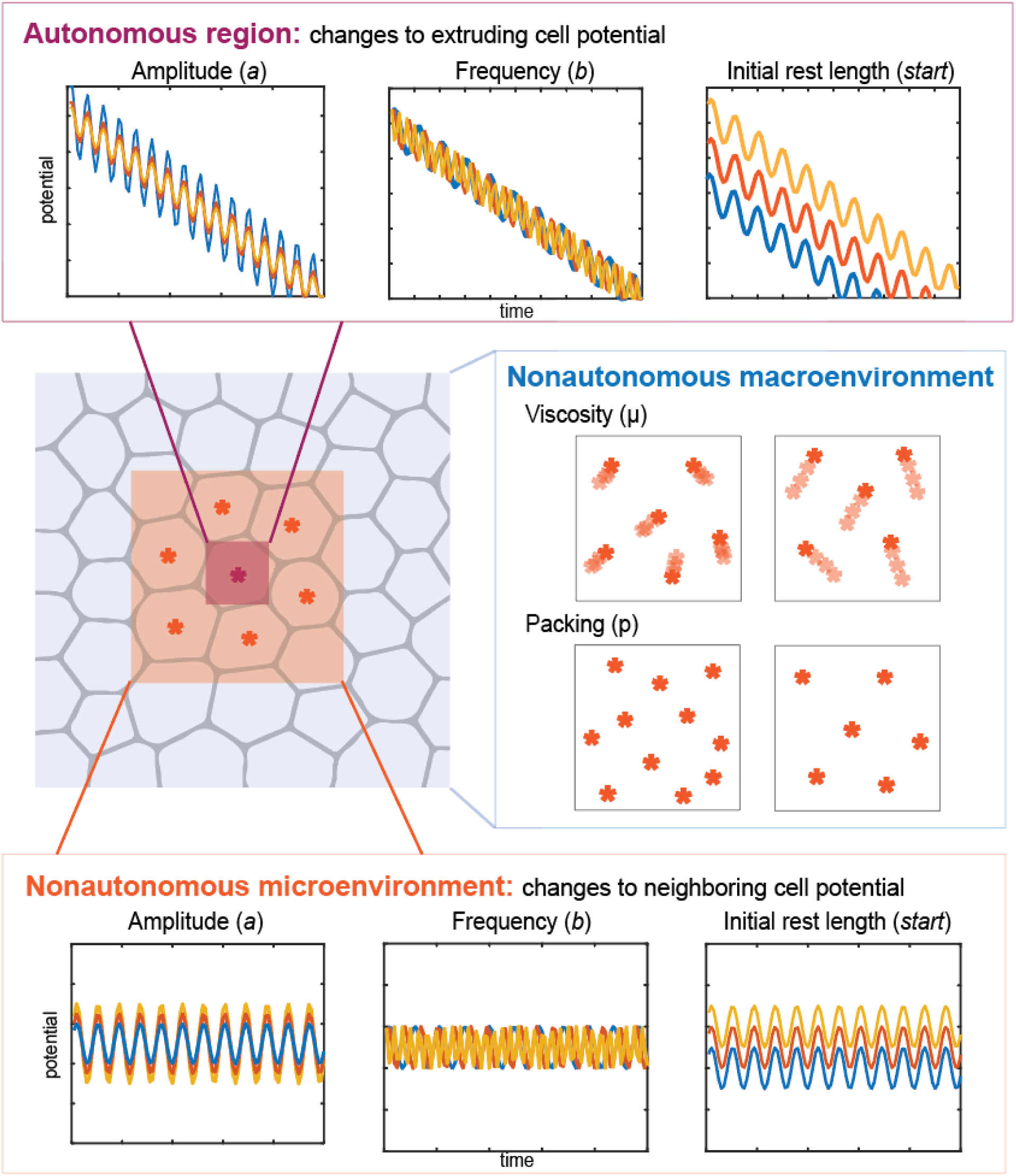
Key parameters used for assessing cell autonomy. Across three scales, the autonomous region, nonautonomous microenvironment, and the nonautonomous macroenvironment, there are eight parameters that are varied to assess their contributions to time to half area loss and the slope of area loss. Asterisks indicate cell centroids and the colors represent different scales, with red being the smallest autonomous scale, orange being the nonautonomous microenvironment, and blue being the nonautonomous macroenvironment.

### Extruding cell potential is a key source of variation in extrusion timing

To explore the role of cell autonomous factors on extrusion, we varied properties of the extruding cell and measure how these parameters contribute to the dynamics of extrusion while maintaining the geometry and mechanical properties of neighboring cells. First, we vary the amplitude and the frequency. Ranges are chosen to capture physiologically observed variation across orders of magnitude. Multiple simulations are run with specifically varied parameter sets. Each simulation generates an extrusion over a time series where cell centers are tessellated to reproduce an epithelial array. Characteristic extrusion features t_half_ and the slope_half_ are extracted from the extruding cell area changes during extrusion (Figure 2e). For the same initial potential, we observe differences in t_half_ as amplitude changes (Figure 4a) but do not observe changes in the slope_half_ (Figure 4b). By varying the frequency of oscillation, we do not observe differences in t_half_ (Figure 4c). However, unlike when we changed amplitude, variation in the frequency of oscillation alters slope_half_, especially at high and low frequencies (Figure 4d).

**Fig 4.**
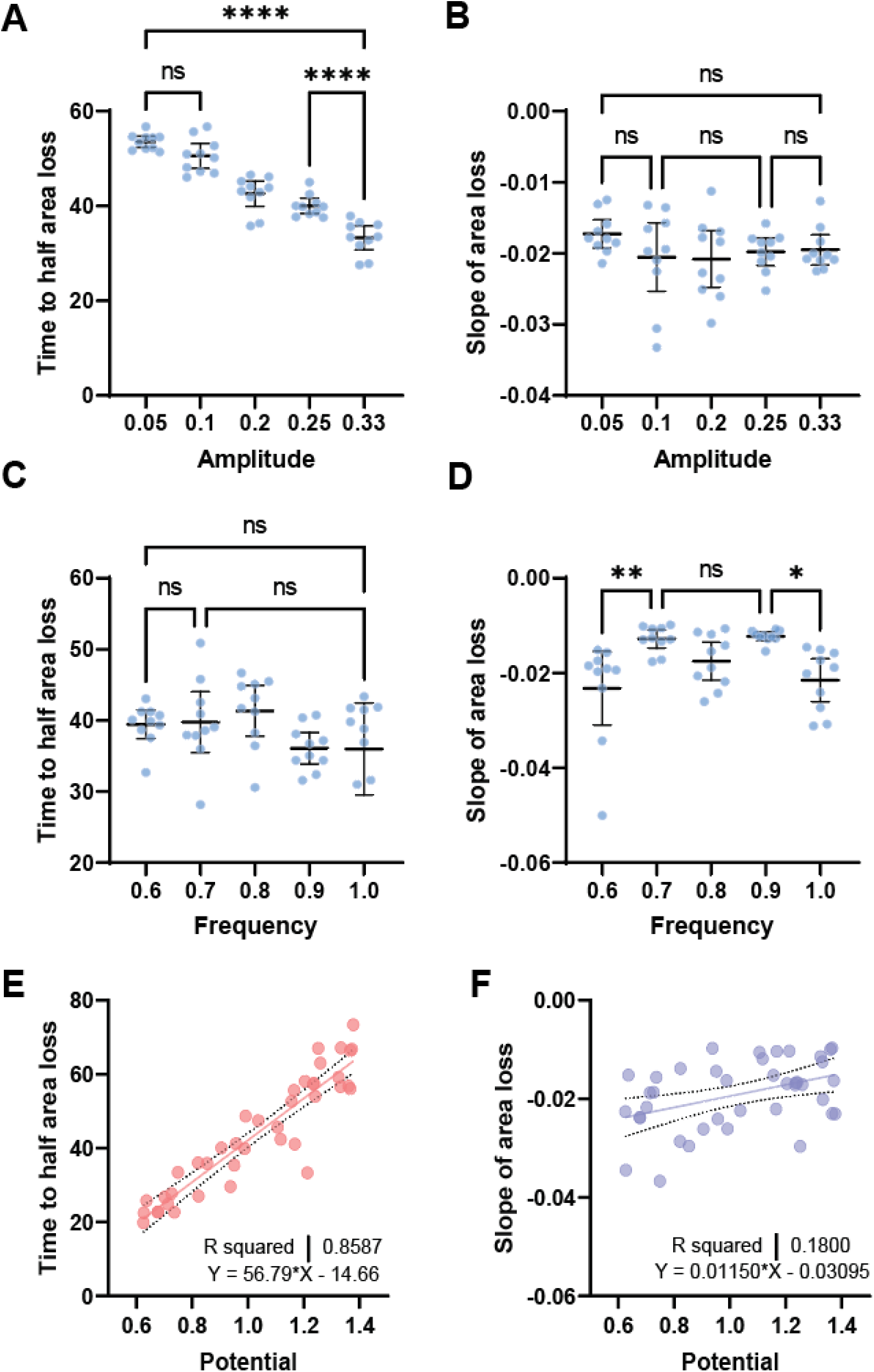
Effects of cell-autonomous properties on extrusion dynamics. (a) Effect of autonomous amplitude change on time on half area loss (b) Effect of autonomous amplitude change on slope of area loss (c) Effect of autonomous oscillatory period change on time on half area loss (d) Effect of autonomous oscillatory period change on slope of area loss (e) initial rest length/potential of the extruding cell as a function of time to half area loss for n=40 simulations (f) slope of area loss related to initial potential of the extruding cell for the same 40 simulations

The initial state or size of a cell may contribute to the dynamics of extrusion. We tested this possibility by using a range by varying the initial potential for extruding cells. The potential of a cell is a major contributor to its size before extrusion. Additional contributions to a cell’s size are the potentials of its immediate neighbors and global packing within the tissue (Figure 1g). From these simulations we find a positive correlation between the initial potential of a cell and the cell’s t_half_ (R^2^= 0.8587) (Figure 4e). Furthermore, slope_half_ has a mild positive correlation with the initial potential of the extruding cell (R^2^ = 0.18) (Figure 4f), indicating cells that are large due to their low potential are also slower to extrude. Observations in zebrafish suggested larger cells might extrude more slowly than smaller cells (Atieh et al., 2021b).

### Cell potentials of neighboring cells and the local mechanical environment can play a role in extrusion timing

Next, we query the contribution of nonautonomous biophysical factors on extrusion dynamics. To isolate the contribution of the mechanical environment of the extruding cell, we hold constant the potential function of the extruding cell and vary the potential of neighbor cells. The neighbor cells’ potential oscillations are set up to vary in a similar way: increasing and decreasing each parameter from their baseline levels of amplitude and frequency. After varying amplitude, we find the amplitude of neighbor cell oscillation has a minimal effect on extruding cell area loss (Figure 5a,b). After varying period, we find little effect on t_half_ and slope_half_ (Figure 5 c,d).

**Fig 5.**
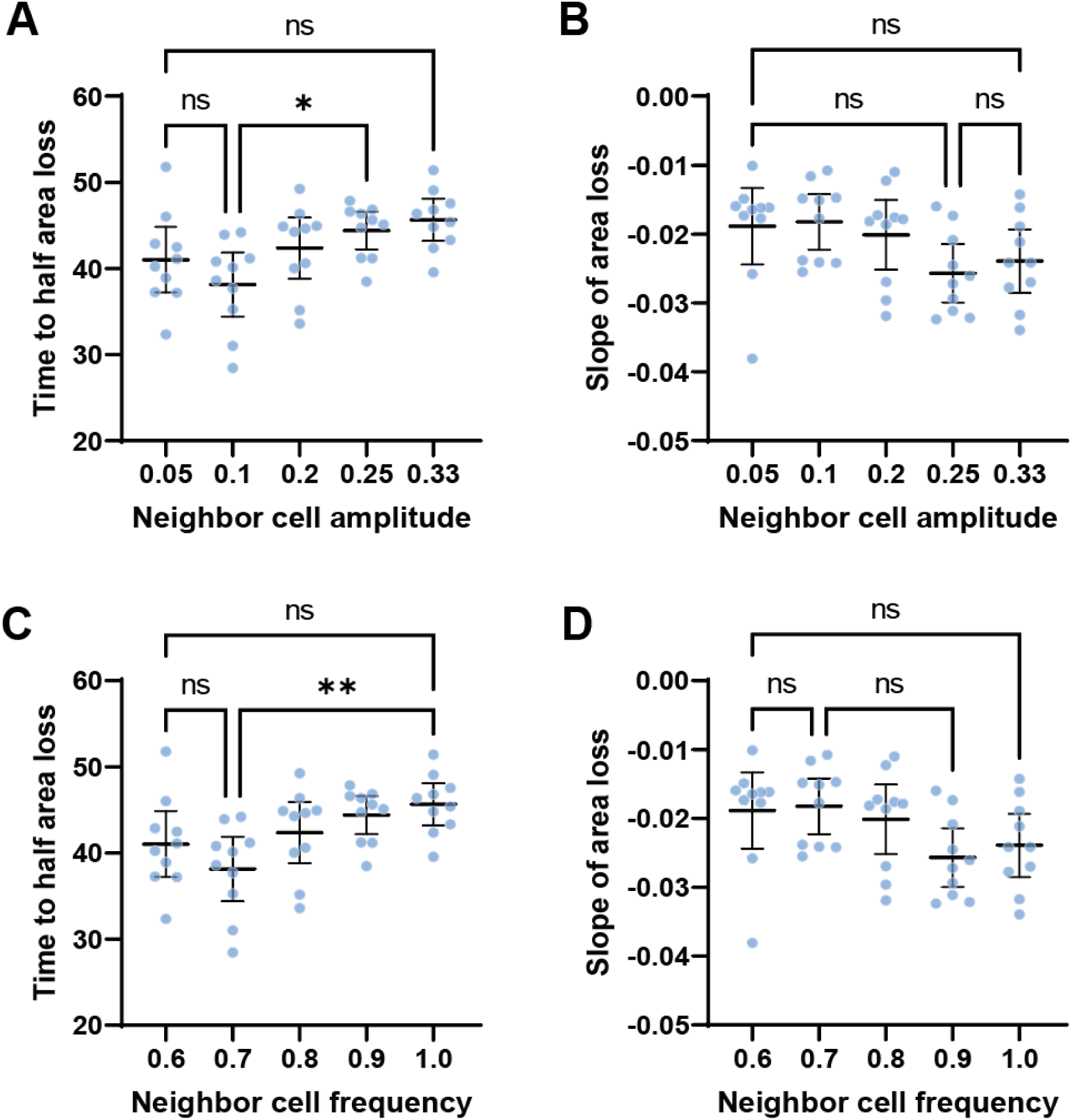
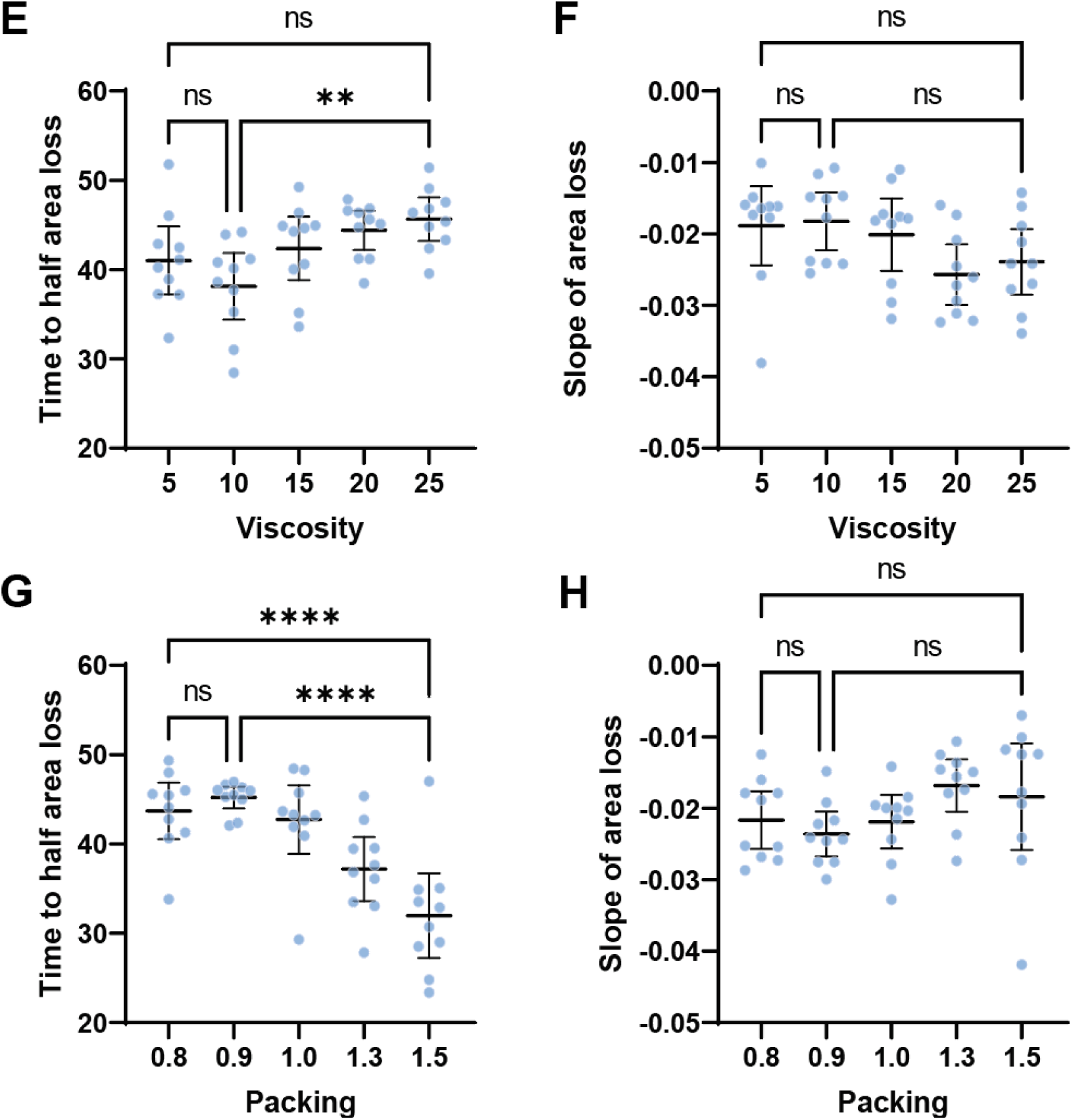
Non autonomous effects from cell neighbor and tissue-scale mechanics. (a) Effect of neighbor cell amplitude potential variation on time to half area loss (b) Effect of neighbor cell amplitude potential variation on slope of area loss (c) Effect of neighbor cell period potential variation on time to half area loss (d) Effect of neighbor cell period potential variation on slope of area loss (e) Effect of viscosity on time to half area loss (f) Effect of viscosity on slope of area loss(g) Effect of packing on time to half area loss (h) Effect of packing on slope of area loss Embedded into figure 6, raw table

After varying the properties of extruding cells or their neighbors we turned to varying the bulk mechanical properties of the tissue, or macroenvironment, including modulus and viscosity. The mechanical status of a tissue may act through mechanobiological feedback to influence the assembly of contractile structures within the cell (Blanchard and Adams, 2011; Chanet et al., 2017); however, we do not include such feedback processes in our model. Viscosity may contribute to extrusion phenotypes due to dampening the effect of forces from contractility-driven oscillations within the tissue (Charrier et al., 2018; Murrell et al., 2011). Changing the viscosity modifies the timescale with which the tissue responds or deforms after an applied force. If contractions from oscillations propagate through the tissue more slowly, this may affect the t_half_ and slope_half_. Changing levels of viscosity have an effect on t_half_ and no effect on the slope_half_ (Figure 5e,f). Another factor impacting the micromechanical environment of the tissue is how cells are confined, or distributed in space, that in turn affects tissue mechanics. Crowding is known to induce extrusion of non-apoptotic (live) cells (Eisenhoffer et al., 2012); therefore, we tested the effect of packing, e.g. cell density, on extrusion dynamics. The prediction is that higher cell density will make cells extrude faster due to having lower areas, similar to the case of low rest lengths (Figure 4e). We find agreement with this prediction, and observe higher levels of packing drives cells to extrude in less time, albeit at a similar rate (Figure 5g,h).

### Sources of variation in extruding cell area loss

Our new model delineates the effects of cell behaviors and tissue mechanics on t_half_ and slope_half_. Yet, due to the stochastic nature of cell behaviors across the field, we were not able to precisely control variation that may contribute to the duration of extrusion. To extract the contributions of stochastic factors and precise contributions to variation in t_half_ and slope_half_, we performed a sensitivity analysis. Previously, when assessing cell autonomous effectors of extrusion, the focus was on the initial mechanical state of the cell and features of its oscillatory resting potential (Figure 4). To determine how these features contribute to variation in outcomes, we sought to consider the initial rest length of the extruding cell and the amplitude and frequency of the rest length oscillation. As non-autonomous factors such as the neighbor cells and mechanical environment also contribute to extrusion, we sought to assess their relative contributions (Figure 5). For neighboring cells, we assessed the parallel features to extruding cells: initial rest length, amplitude, and frequency of oscillation. For the tissue mechanical environment, we again consider viscosity and packing, as they appeared to contribute to extrusion dynamics (Figure 5e-h).

To define the ranges of the eight model parameters and make comparisons relative, we matched ranges for parameters of the extruding and the neighboring cells. We tested the contribution of a range of initial rest lengths to the geometric heterogeneity of the model epithelium (Figure 1f-h). Given the impact of rest length on area and sidedness (Figure 1h-i) and the wide range of areas and sidedness seen *in vivo* (Figure 1c-d), we selected cell potentials from the high-width uniform distribution. For the amplitudes and frequencies of the extruding cells and their neighbors, the same ranges/deviations from the base case were used. For determining the ranges of viscosity and packing in the model, direct literature comparisons were lacking, so we similarly chose values greater and lesser than that of the experimentally determined base case to capture variation. Simulations carried out at the bounds of parameter ranges were visually inspected for stability of the tessellation and segmentation.

To assess parameter contributions to extrusion dynamics, we used extended Fourier Amplitude Sensitivity Testing (eFAST; Saltelli et al., 1999). To implement the eFAST technique, we sample input parameters from periodic distributions. Associated model outcomes and peaks thereof are used to calculate Sobol indices to quantify parameter contributions to time to half area loss and the slope of the area loss (t_half_ and slope_half_). Two Sobol indices are calculated (Figure 6a); the first order index (S1) reports how sensitive the output is to a parameter’s variation, with high values indicating more sensitivity; the total order index (ST) reports synergistic effects of any order. Confidence intervals of these indices are also provided (Table S1).

**Fig 6.**
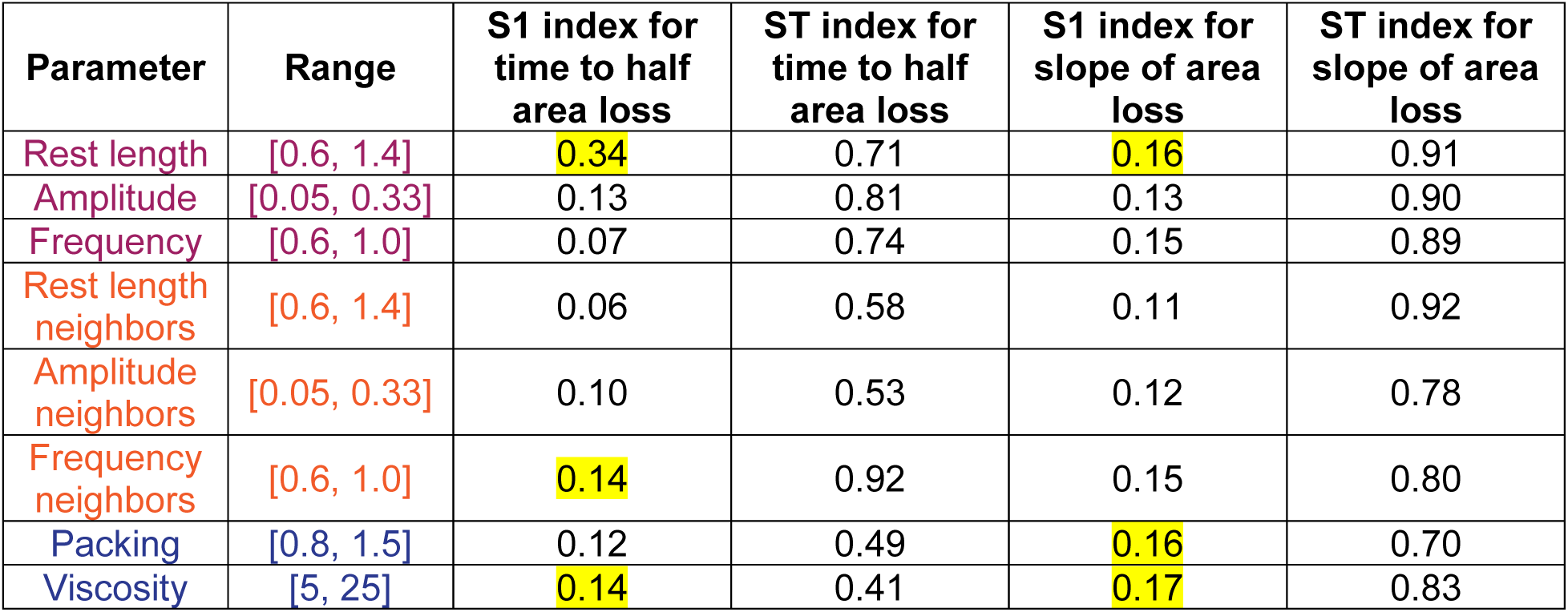
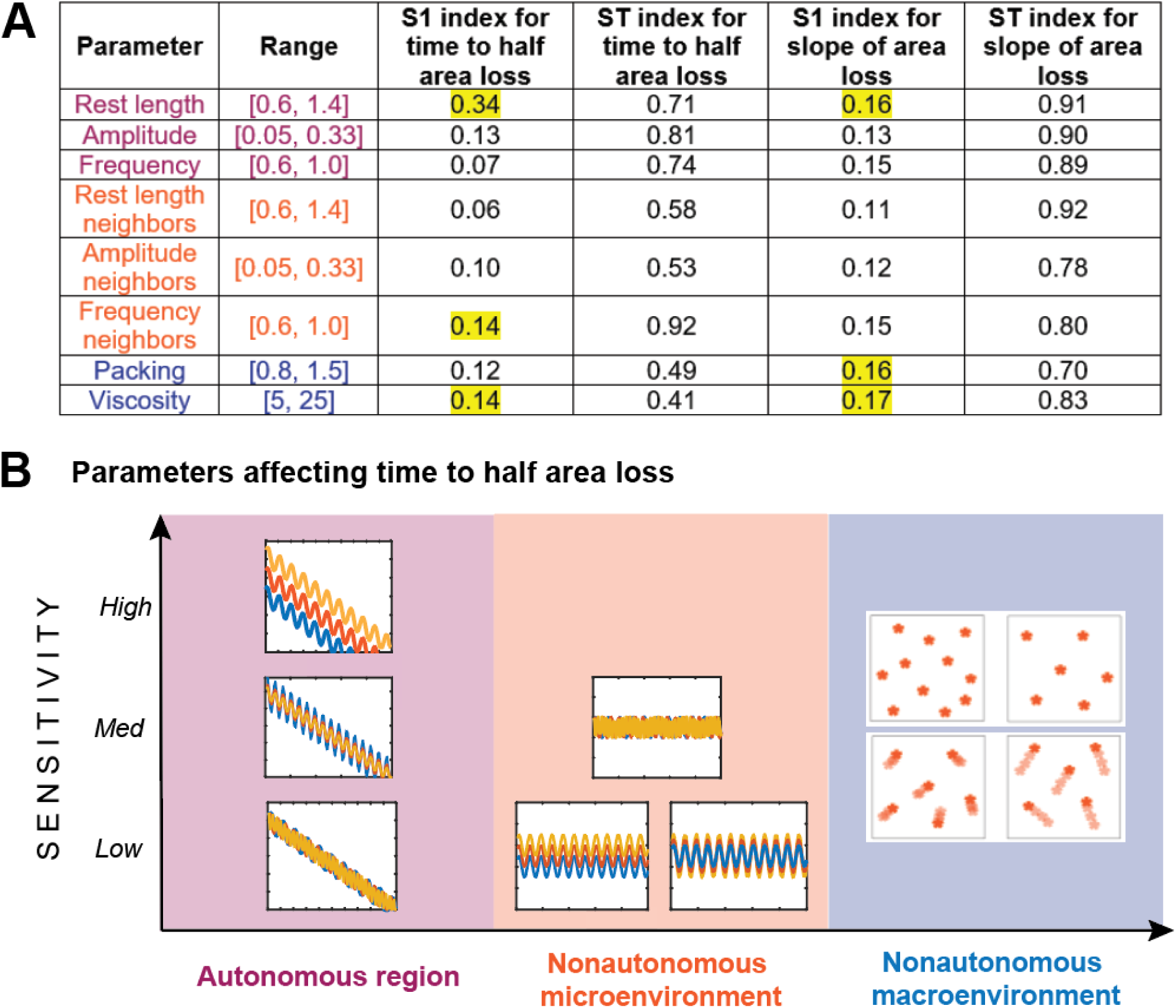
Summary of contributors to extrusion dynamics. (a) Table with parameter ranges for sensitivity analysis, with first (S1) and total (ST) order indices for time to half area loss and slope of area loss. Highlighted values represent the top 3 contributors to S1 indices/sensitivity. (b) Parameters affecting time to half area loss sorted by scale and level of contribution by S1 Sobol index (low= 0-0.1, med= 0.1-0.3, high= 0.3+). Colors of text in table correspond to different scales introduced in figure 3, with the autonomous region being the smallest in red, nonautonomous micro- and macro-environments in orange and blue, respectively.

Model parameters with the greatest effect on t_half_ are the extruding cell’s initial rest length, amplitude, frequency of the neighbor extruding cell potential oscillation, and tissue viscosity. For all of these, the total Sobol index exceeds S1, indicating high order interactions between multiple parameters are affecting the outcome. Neighbor cell potential features have a lower impact on synergistic effects. For slope_half_, the parameters contributing to the most variance are the rest length of the extruding cell, the packing of the tissue, and viscosity. The S1 index is close among all parameters, with neighbor cells having slightly lower values for both indices. This, in the context of our other results, indicates that smaller cells that are pulsing more strongly, with faster pulsing neighbors, moving more quickly, and more confined in the tissue will extrude at a faster rate. By binning the S1 Sobol indices across scales, we are able to gain insight into the role of individual parameters on the entire process of extrusion (Figure 6b). Both t_half_ and slope_half_ are affected more strongly by the initial amplitude of the potential of the extruding cell and its rest length of the extruding cell and tissue viscosity. The frequency of the extruding cell’s potential has a minimal effect on the t_half_, but a strong effect on slope_half_ compared to other parameters, indicating that the number of contractions affects extrusion dynamics differently than the strength of the contractions. Thus, nonautonomous properties of the microenvironment have only a moderate impact on extrusion dynamics.

## DISCUSSION

Our computational model of oscillatory cell extrusion can represent dynamics in a way that enables precise perturbations that cannot be made *in vivo*. Properties of the repulsive potential function of the extruding cell largely determine the morphogenesis of extruding cells, including time to half area loss (t_half_) and slope of area loss (slope_half_). Furthermore, tissue wide properties including viscosity and packing enhance cell extrusion. Using our model, we observe how extrusion changes as non-autonomous properties, such as neighboring cell mechanics are altered, even as we keep the extruding cell properties fixed. In contrast to factors regulating the behaviors of neighboring cells, the biophysical properties and mechanics of the extruding cell have large effects on t_half_ and slope_half_. Our comparative analysis of model parameters and constraints suggest that biophysical features of the extruding cell such as the potential function and rest length, as well as tissue wide viscosity and packing contribute the most to t_half_ and slope_half_. In particular, we find that smaller cells with higher amplitude area oscillations that are more confined or in a more viscous environment will be removed from the tissue faster than other cells.

Active particle modeling together with hybrid meshing enables variation of mechanical properties and observation of their impact on the homeostasis of epithelial cell sheets. Here we focused on the mechanics of extrusion from the perspective of the extruding cell and its mechanical environment. We are able to represent a diverse apical morphology of cells within an epithelium and manipulate both an extruding cell and its neighbors. With a common analysis pipeline, applied to both live and simulated cell timelapses, we are able to make comparisons between these cases that were not previously possible. This type of model can be used to represent processes in a dynamic tissue, as opposed to more traditional models focused on static tissues with isodiametric cells. Existing traditional vertex models including the extrusion process do not include an explicit extrusion program (Marinari et al., 2012b) and don’t rigorously quantify differences in contributions of the extruding cells and their neighbors. Our new model addresses these limitations to provide insights into extrusion programs under different conditions while echoing modeling findings that properties of the extruding cell itself can be a large contributor to extrusion dynamics (Kuipers et al., 2014).

Using this new model, we find autonomous contraction is a key first step to promoting extrusion of cells from epithelial tissues. These data are consistent with the apoptotic cell being an active physical participant in the process of cell extrusion (Kuipers et al., 2014). During extrusion, an actomyosin ring forms in the extruding cell (Villars and Levayer, 2022) and pulses of contractile medio-apical actomyosin drive cell-autonomous constriction that contributes to cell expulsion (Atieh et al., 2021b). This contractility and rosette formation has been proposed to provide an essential mechanical signal to the surrounding cells by transiently increasing junctional tension and local cellular density (Kuipers et al., 2014). Our results provide additional support to the idea that the initial size/state of the cell may dictate the response to the environment (crowding) or damage (in this case, MTZ). Defining the autonomous features that promote extrusion could aid in predictions of cells primed to leave the tissue after a stimulus or during pathogenesis.

In our model, extrusion is not an emergent process—it is encoded in the extruding cell’s potential. Elucidating factors triggering extrusion *in vivo* can drive the development of future models where specific cell-intrinsic factors result in the emergence of extrusion without explicit instructions. What are the molecular signals that drive this autonomous contraction program and extrusion in response to different stimuli? During apoptotic cell extrusion *in vivo*, caspase activation leads to enrichment of the bioactive lipid sphingosine-1-phosphate that regulates pulses to dictate regions of extrusion (Atieh et al., 2021b). Caspase activation has also been shown to promote the disassembly of microtubules that is required for progression of extrusion (Valon et al., 2021). Further, a wave of calcium emanating from the apoptotic cell can induce polarized movement of the surrounding cells toward the extruding cells (Takeuchi et al., 2020). A current limitation of our computational model is the incorporation of molecular components that dictate specific phases of the extrusion process. Integration of known molecular regulatory components of actomyosin assembly/disassembly cycles that drive oscillatory dynamics into this new active particle-based model will provide key insights into removal of defective cells from homeostatic epithelial tissue maintenance.

While our data are supportive of oscillatory dynamics of an extruding cell providing the strongest contribution to the trajectory of extrusion, the neighboring cells were predicted to have a less pronounced contribution. Existing models representing contraction (without apoptosis) in other model systems note the strength of contractions as a pertinent parameter (Cavanaugh et al., 2020; Clement et al., 2017). Contraction of the apoptotic cell also triggers a coordinated elongation of the neighboring cells, which requires E-cadherin-mediated cell-cell adhesion (Lubkov and Bar-Sagi, 2014). The small GTPase RhoA is a cadherin-dependent signal in the neighbor cells that becomes activated in response to contractile tension from the apoptotic cell (Duszyc et al., 2021). However, in this study, changes in cell adhesion within the extruding cell or its neighbors were not directly accounted for. Therefore, it is plausible that the neighbors play a more significant role in facilitating the exit of unwanted cells than reflected by our model. Future studies using optogenetic manipulations of caspase activation, contractility, and adhesion in the extruding cell and its neighbors prior to elimination will provide additional clues on the mechanics of cell elimination during tissue homeostasis.

Lastly, our model also suggests that radial intercalation, or the insertion of a cell into a stable epithelium, is not simply the reverse of cell extrusion. The precise biophysical events driving cells to migrate from a basal population toward the apical surface and insert are beginning to be elucidated (Rao and Kulkarni, 2021; Stubbs et al., 2006; Ventura and Sedzinski, 2022). At the conclusion of extrusion, the extruding cell is surrounded by a high order array of cell neighbors present from to the start of extrusion. By contrast, radial intercalating cells first appear at tri-cellular junctions. While extrusion may seem analogous to the process of radial intercalation, the assembly of apical junctions in radial intercalation involves distinctive biophysics and molecular complexes than their disassembly during extrusion. A Cellular Potts model of radial intercalation from a quasi-2D perspective (Szabó et al., 2016) shows the inverse process in theory, but not in practice. A finite element model showing radial intercalation has intercalations driven by protrusive activity and a tension gradient, with intercalation events being randomly selected (Neumann et al., 2018). While we do not represent this polarity explicitly given the two-dimensional nature of our model, the phenomenology of apical area loss aligns with what is observed in the larval zebrafish epidermis.

In summary, our new computational model based on *in vivo* observations provides a new tool to assess the contributions of the extruding cell and its neighbors to the process of eliminating defective cells from epithelial tissues. Given the conserved nature of epithelial topology and extrusion in different organisms, our model could provide insights into additional mechanisms and guide future investigations into regulation of epithelial tissue homeostasis.

## METHODS

### Zebrafish

Experiments were conducted on larval zebrafish (*Danio rerio*) maintained under standard laboratory conditions with a cycle of 14 h of light and 10 h of darkness. Larvae were collected and kept in E3 larva medium at 28.5°C and staged as previously described (Westerfield, 2007). The zebrafish used in this study were handled in accordance with the guidelines of the University of Texas MD Anderson Cancer Center Institutional Animal Care and Use Committee.

### Induction of extrusion in surface epithelial cells in larval Zebrafish

4dpf larvae were treated with 10mM metronidazole (MTZ) for 2-4 hours in a 28.5°C incubator to induce extrusion activity, as described in (Atieh et al., 2021a; Atieh et al., 2021b).

### Imaging of Zebrafish larvae

4dpf zebrafish larvae were anesthetized with 0.04% tricaine in E3 and mounted in an X-plate petri dish with using 0.5% UltraPure Low Melting Point Agarose and E3. The larvae were imaged on a ZEISS LSM 800 laser scanning confocal microscope with fluorescent channels set to a wavelength of 488nm and 561nm. Images were acquired from 30sec, 1 minute or 1 minute and 30 second acquisition times.

### Quantification of Zebrafish cell area and polygon topology

The resulting confocal images were handled in Fiji and segmented and the resulting area and polygon topology were assessed using Tissue Analyzer (Etournay et al., 2016) as described in the previous protocols paper (Atieh et al., 2021b).

### Simulation and analysis pipeline

The simulation is coded as an ImageJ macro to allow pixel manipulation and visualization. The simulation is performed, and centroids are tessellated using a custom plugin implementing the hybrid CVT. Segmentation of the cells is performed via ImageJ using cell centroid information derived from the model and assigning unique pixel values to cells detected as particles after tessellation. We extract morphometric features using regions of interest (ROIs) from each frame of the simulation (typically 60-70 frames). Sidedness is calculated for cells not on the boundaries of the simulation by expanding the ROI and finding unique pixel intensities. We calculate strain from ROI measurements for cells over time using MATLAB (Mathworks). To calculate the slope and time to half area loss, we calculate the lower Hilbert envelope by using the discrete Fourier transform on the analytic signal of the strain data. The envelope in the neighborhood of half area loss is used to calculate the slope and interpolate the time at which half area loss is achieved. Statistical analysis is performed in MATLAB and GraphPad Prism 9. To speed up the simulation and analysis pipeline, we used MATLAB-ImageJ (Sage et al., 2012). By running ImageJ via MATLAB, simulations could be run in sequence without jumping from MATLAB to ImageJ between simulation and analysis phases. The simulation and analysis pipeline overview is provided (Figure S2).

### Statistical Methods

To extract amplitudes from the area strain trajectories, we calculate the Hilbert envelope by using the discrete Fourier transform on the analytic signal of the strain data. Amplitude is calculated as half the difference between adjacent peaks and troughs in the signal for live cell and simulated data. For comparisons between two groups such as the amplitudes for matching the base case, we used the Wilcoxon rank-sum test at the 95% significance level. For analysis of manipulated features such as amplitude, frequency, and packing, we use a one-way ANOVA to compare variations across groups at the 95% significance level. We checked normality of the distribution of half times as well as the residuals, and our data meets criteria for sufficient samples to account for deviations from normality.

For the attribution of features of time to half area loss, we performed global sensitivity analysis using the extended Fourier Amplitude Sensitivity test (eFAST) using the SALib package in Python (Herman and Usher, 2017). Results are interpreted as contributions to variability in the output variables of time to half area loss and slope of area loss, with larger values indicating a greater contribution.

## Acknowledgements

We would like to thank members from the Eisenhoffer and Davidson labs for their support and helpful discussions. This work was supported by grants from the National Institutes of Health, R01GM124043 and R35GM149226 to GTE, and R01 HD044750, R37 HD044750, and R21 HD106629 to LAD. Additionally, SA was supported by the Biomechanics in Regenerative Medicine (BiRM) Training Grant from the NIBIB (T32 EB003392).

## Author contributions

SA: Conceptualization, Software, Writing-Original Draft; LT: Investigation, Validation; YA: Methodology, Conceptualization; GTE: Conceptualization, Writing-Review & Editing, Supervision, Resources; LD: Conceptualization, Software, Writing-Review & Editing, Project Administration, Methodology.

## Lead contacts

Further information and requests for resources and reagents should be directed to the Lead Contacts: Lance Davidson and George Eisenhoffer.

## Materials availability

The transgenic zebrafish lines used in this paper are available from the George Eisenhoffer upon request.

## Data and code availability

Data supporting the findings presented can be found within the body of the paper and the supplementary text and videos. Data supporting the findings presented can be found within the body of the paper and the supplementary text and videos. Additional information and code for the computational model have been depositied in a Dataverse repository-

Anjum, Sommer, 2023, "Data for "Assessing mechanical agency during apical apoptotic cell extrusion"", https://doi.org/10.7910/DVN/XKVWGA, Harvard Dataverse, DRAFT VERSION Additional information on the data and computational model can be requested from the corresponding authors.

## Notes

### Competing Interest Statement

The authors have declared no competing interest.

