## Supplementary Information for "Assessing mechanical agency during apical apoptotic cell extrusion"

**SUPPLEMENTARY INFORMATION for Anjum et al. (submitted).**

CONTENTS

*Model description*

*A hybrid tessellation technique capable of representing oscillatory cell extrusion*

*Low local strain occurs in cases of extrusion*

*A larger field model with distributed extruding cells is computationally expensive, but mimics in-vivo cell networks*

*Extended Fourier Amplitude Sensitivity Test (eFAST) global sensitivity analysis*

*Figure S1, Supporting Fig 2. Schematic of hybrid tessellation scheme*

*Figure S2. Simulation and analysis pipeline steps*

*Figure S3. Different packing scenarios and their contribution to cell morphology to determine a packing regime*

*Table S1. Global sensitivity analysis continued*

*Figure S4. Candidate metrics for describing cellular and tissue deformations*

*Figure S5. Simulation of a larger cell field*

*Supplementary Video 1. Cell area oscillations* in vivo

*Supplementary Video 2. Simulation with extruding cell experiencing high amplitude oscillations*

*Supplementary Video 3. Simulation with extruding cell experiencing low amplitude oscillations*

*Supplementary Video 4. Simulation with extruding cell experiencing high frequency oscillations*

*Supplementary Video 5. Simulation with extruding cell experiencing low frequency oscillations*

*Supplementary Video 6. Simulation with neighboring cells experiencing high amplitude oscillations*

*Supplementary Video 7. Simulation with neighboring cells experiencing high frequency oscillations*

*Supplementary Video 8. Simulation of a larger field with five extruding cells*

#### *Model description*

We have developed a lattice-free model of an epithelial sheet. Cells are described as nodes that are initialized randomly within a square bounding box. Corner nodes are fixed, while border nodes move along one axis, and interior nodes may move along both axes. Nodes interact with each other by repelling neighbors via a nonlinear compressive spring maintaining cell areas. Cell movement is governed by a force balance enforcing Langevin dynamics, incorporating stochasticity and drag forces from viscous media. At each timestep, centroid positions are advanced using forward Euler methods. At the beginning of the simulation, cells are randomly assigned uniformly distributed potentials/rest lengths. These rest lengths determine how much cells repel neighboring cells, as a function of distance. The repulsion force is scaled by viscosity μ and the simulation is advanced using forward Euler methods. The cell centroids are then tessellated using a custom modified Voronoi tessellation implemented by a java-based FIJI plugin. When cells are first initialized, they move more because of their proximity causing more force due to their random placement. We allow 100 timesteps for the cell positions to equilibrate before performing manipulations. An extruding cell is selected from centroids in the center of the field at random, and centroids within a pixel based radius are selected to be the neighboring cells and assigned an oscillatory potential as well.

#### *A hybrid tessellation technique capable of representing oscillatory cell extrusion*

We developed a new method of tessellating cells that represents the physics of the system better than a Voronoi tessellation. The Voronoi tessellation is done purely based on the location of the cell centroids. First, a Delaunay triangulation is performed, and the circumcenters of these triangles are connected to form the Voronoi tessellation. In our new method, we calculate the centroid of the triangle, weighted by the resting lengths of the cells that form the triangle. We average this with the original Voronoi circumcenter. This enables us to represent cellular extrusion by dramatically lowering a cell's resting length in the simulation and inflating this value during the tessellation to bias the weighted centroid toward an extruding cell. A shortcoming is sudden T1-type flips of edges in the network that occur when the weighted circumcenter moves outside of the Delaunay triangle. We can attribute this to the special case in which the circumcenter of the Delaunay triangle lies outside the triangle itself and when Delaunay triangles undergo large configuration changes. Due to this instability, we sought to find the best of both worlds. We tested weighted combinations of the Voronoi tessellation (VT) and the Weighted Centroidal Voronoi (weighted CVT) tessellation. The combination that allowed for the greatest stability while still being able to satisfy our base case criteria was an even (50% each) weighting of the Voronoi tessellation and weighted CVT, which we call the Hybrid Centroidal Voronoi tessellation (hybrid CVT). This new method allows removal of cells in a way that can be precisely controlled.

$$x_{centroid} = \frac{x_i/rest_i + x_j/rest_j + x_k/rest_k}{1/rest_i + 1/rest_j \ + 1/rest_k}$$

This tessellation is implemented via modification of the Delaunay Voronoi ImageJ plugin.

***Low local strain occurs in cases of extrusion***

During exploratory simulations of oscillatory extrusion, strain of a local neighborhood of cells was examined in addition to cell strain. The deformation gradient matrix F is derived from the equation of an ellipse fitted to the ROI of a cell or its neighborhood and is used to calculate Lagrangian finite strains.

$$\varepsilon = \frac{1}{2}(F^T F - I) = \begin{bmatrix} \varepsilon_{xx} & \varepsilon_{xy} \\ \varepsilon_{yx} & \varepsilon_{yy} \end{bmatrix}$$

The x and y directions of cell strains were essentially equivalent, so we chose to show only area strain for clarity in the paper. The local strain of a cell, its neighbors, and its neighbor's neighbors (2-level corona) was negligible, even in the case where only the extruding cell was oscillation, even though it reached half area loss sooner (Figure S4 a,c,d). We expect that this is due to area conservation: when a cell has a shrinking area, the neighboring cell takes on that area. This is not a "perfect 1:1" area allocation as viscous drag forces delay area allocations as a function of force dampening within the media. This is an interesting observation in that extruding cell oscillations may have a varying effect on the deformation of nearby cells. In conjunction with possible minimal autonomous oscillations of non-extruding cells, oscillations may be triggered later on with compounding of these forces.

***A larger field model with distributed extruding cells is computationally expensive, but mimics in-vivo cell networks***

The results shown in this paper involve a model representing a single cell oscillating in a small field of cells to isolate the effects of the mechanical properties studied and establish case-by-case independence on our statistical analysis. However, in-vivo, multiple cells oscillate within a larger field. Another version of the model involves a larger field of approximately 400 cells wherein five cells are designated as oscillatory extruding cells (Figure S5). The extruding cells are within the center third of the field to minimize boundary effects. This adds a layer of complexity, as oscillations from one extruding cell, in close enough proximity to the neighborhood of another extruding cell, may impact the dynamics of that extruding cell, as well as the morphology of its neighbors. This extended model is more computationally expensive but allows a way to probe tissue-scale mechanics of propagating oscillations more directly.

#### *Extended Fourier Amplitude Sensitivity Test (eFAST) global sensitivity analysis*

The eFAST test leverages periodic sampling within the parameter intervals to detect peaks in the output data that are attributed to these signals by computing the Fourier transform. This informs calculation of Sobol indices by measuring the fraction of the variance in the output attributed to Fourier coefficients associated with each of the varied parameters. We used 480 parameter sets generated by the SA Lib fast sampler and ran as many simulations to extract our two outputs. These indices tell us the fraction by which the variation of the summary statistic would be reduced if a parameter was held constant. Together, the effects of synergy between changing multiple parameters are shown. The result for time to half area loss and slope of area loss are very similar, as similar parameters have similar effects on the outcomes.

### ***SUPPLEMENTARY FIGURES***

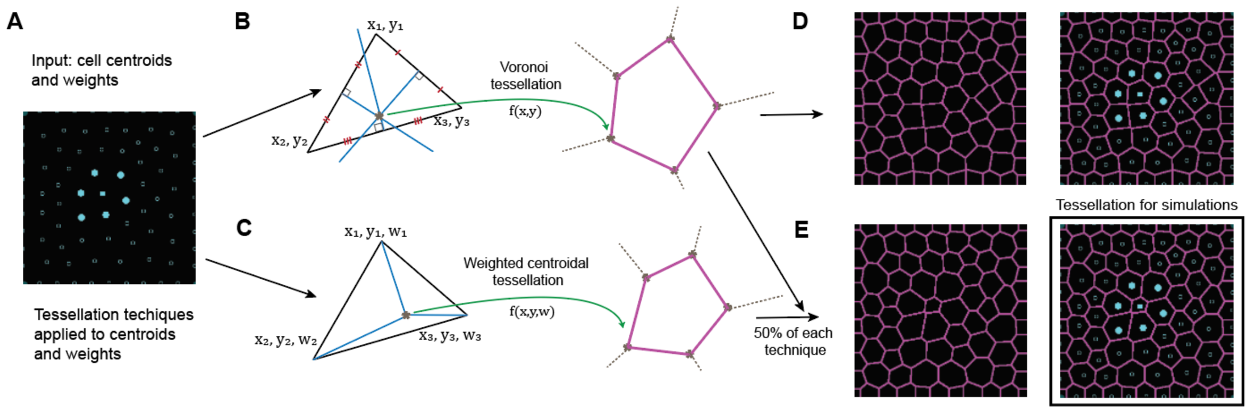


**Figure S1,** Supporting Fig 2. *Schematic of hybrid tessellation scheme* (a) inputs to the tessellation scheme (b) Voronoi schematic using the intersection of circumcenters i.e. purely geometric information (i.e. x and y) (c) centroidal schematic using x,y, and weights corresponding the potential of a centroid, incorporating physical information (d) the Voronoi tessellation of an extruding cell (e) the hybrid tessellation combining B and C such that the cell extruding looks like it undergoes a substantial loss in area, unlike the purely geometric case in D

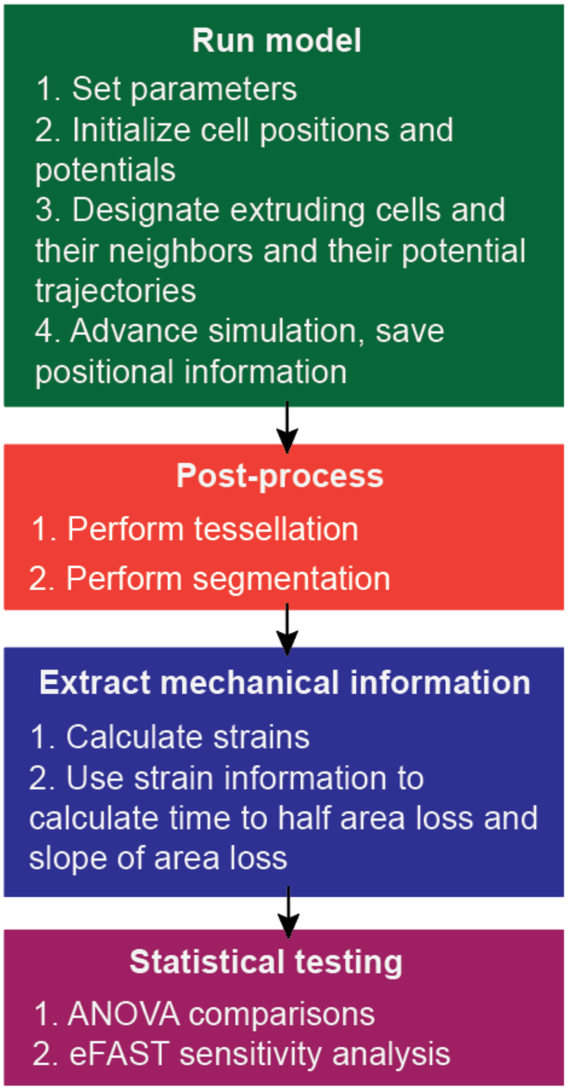


**Figure S2.** *Simulation and analysis pipeline steps.* Simulation step of pipeline with parameter setting, model run steps, and tessellation of cell centroids. Analysis step of pipeline with preprocessing segmentation, followed by calculation of mechanically relevant quantities, and described statistical analyses.

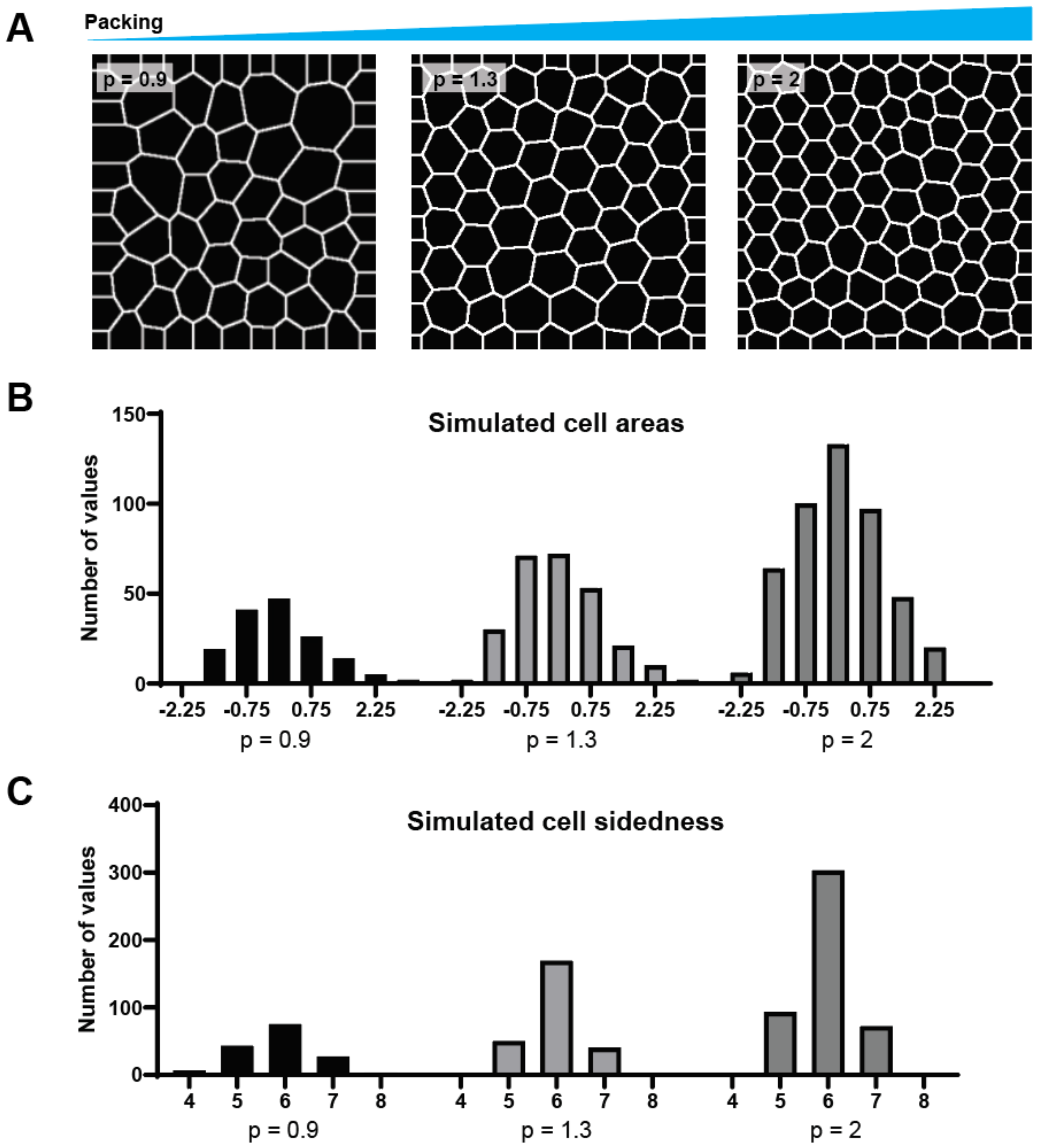


**Figure S3.** *Different packing scenarios and their contribution to cell morphology to determine a packing regime* (a) Representative simulations of packing of cells for different levels of packing factor n using a low-width uniform distribution of potentials (b) Z-score normalized cell area distributions for the three packing scenarios over 10 simulations each (n=154,261,469) (c) Cell sidedness distributions for the three packing scenarios over 10 simulations each (n=154,261,469)

**Table S1.** *Global sensitivity analysis continued*

| Parameter | S1 confidence for time to half area loss | ST confidence for time to half area loss | S1 confidence for slope of area loss | ST confidence for slope of area loss |
|---|---|---|---|---|
| Rest length | 0.22 | 0.14 | 0.18 | 0.08 |
| Amplitude | 0.21 | 0.13 | 0.24 | 0.15 |
| Frequency | 0.23 | 0.11 | 0.16 | 0.12 |
| Rest length neighbors | 0.24 | 0.11 | 0.20 | 0.11 |
| Amplitude neighbors | 0.23 | 0.15 | 0.24 | 0.11 |
| Frequency neighbors | 0.22 | 0.13 | 0.27 | 0.12 |
| Packing | 0.21 | 0.13 | 0.23 | 0.10 |
| Viscosity | 0.20 | 0.12 | 0.24 | 0.12 |

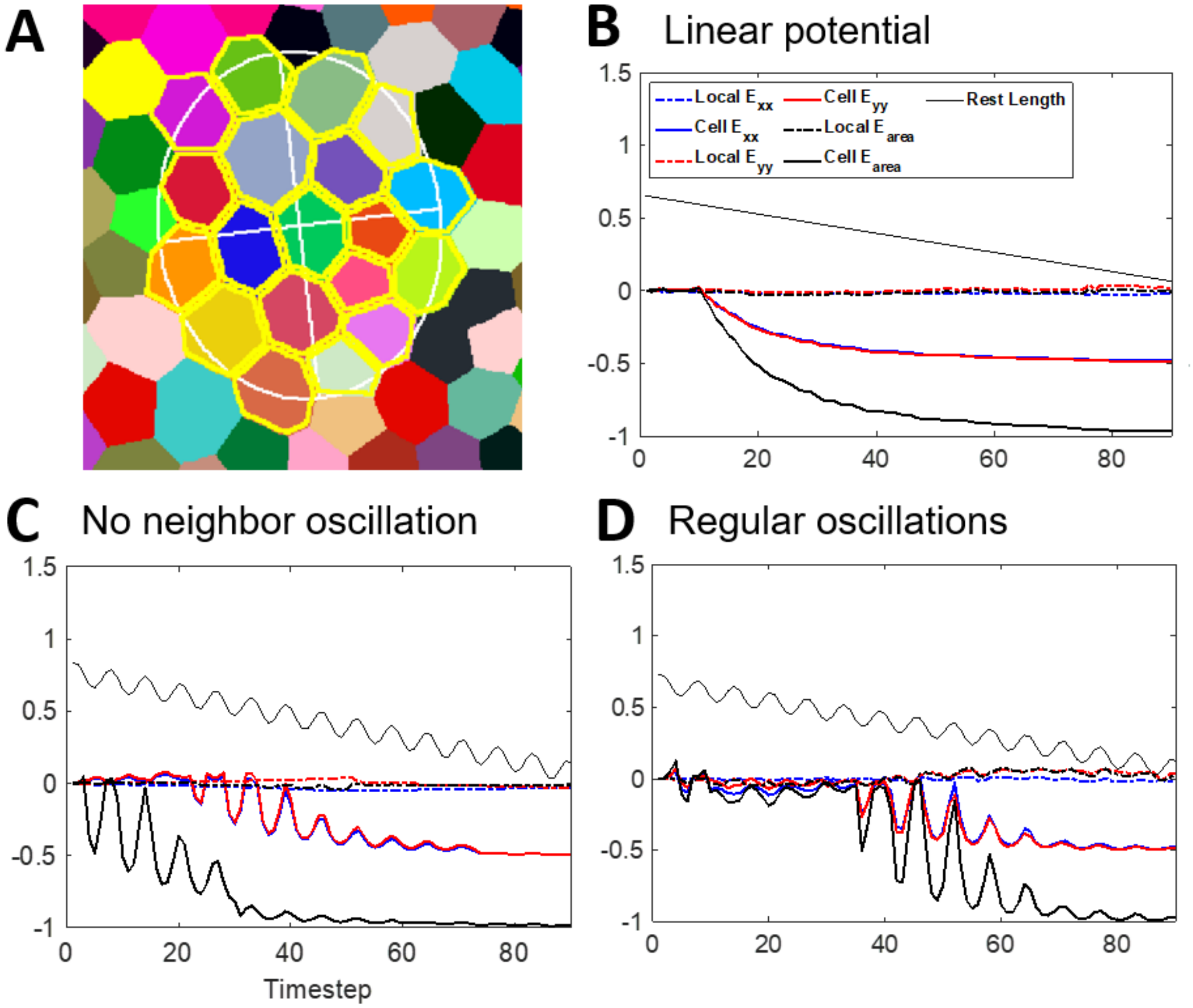


**Figure S4.** *Candidate metrics for describing cellular and tissue deformations* (a) Designation of a two-level corona of a cell, its neighbors, and its neighbors' neighbors. An ellipse is fitted to this domain and its deformation is used to calculate local strain. (b) Trajectory of cellular and local strains of a cell with a linearly decreasing potential, with no oscillations following a trajectory similar to oscillatory extruding cells. (c) Strains of a cell oscillating without neighbor oscillations like in the body of the paper. (d) Strains of a cell with neighbor oscillations, as shown in the body of the paper.

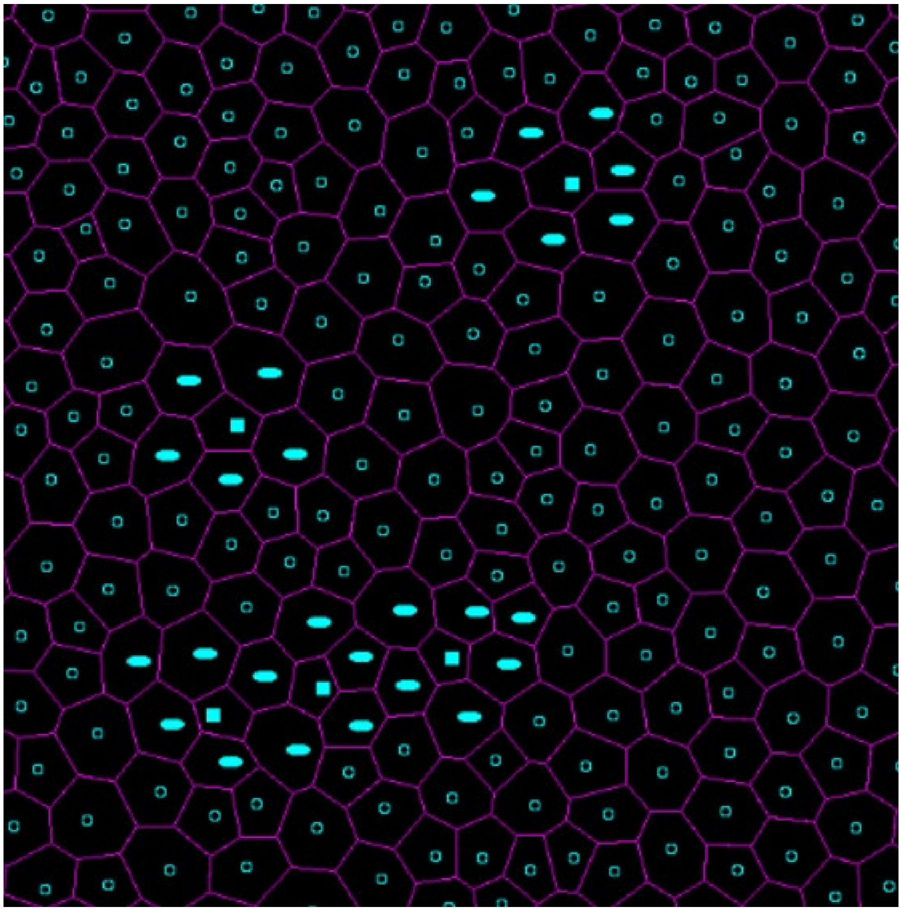

**Figure S5.** *Simulation of a larger cell field* (a) Much larger field of cells in which 5 cells are randomly selected to be extruding cells, denoted with filled in squares, with oscillating neighbors shown as ovals. This better mimics the *in vivo* situation where extruding cells can be near or far from each other. The same simulation and analysis steps can be used on this type of simulation, at great computational cost.
